# Concept Learning Builds Behaviourally Relevant Attentional Templates

**DOI:** 10.1101/2025.09.29.679121

**Authors:** Melisa Gumus, Zoey Zhi Yi Lee, Michael L. Mack

## Abstract

Attention optimizes learning by filtering relevant information to build conceptual knowledge. However, how learned concepts, once encoded in memory, subsequently guide attentional processes remains an intriguing question. We propose that concept learning leads to the emergence of attentional templates that store goal-relevant representations, thereby actively guiding attention allocation. Participants completed two separate learning tasks and a test, wherein each trial began with a cue, indicating which learning task should be employed. Random test trials included a probe instead of concept specific features: a small arrow appeared at a feature location that was relevant (i.e., valid) or irrelevant (i.e., invalid) for the cued task.

Successful learners were faster at responding to valid probes than invalid, demonstrating the deployment of concept-specific attentional templates. Importantly, the efficiency of this attention allocation was tied to concept learning success, with higher learning performance yielding greater response time benefits at test. Thus, our results reveal that learning builds behaviourally relevant attentional templates, and subsequently, learned concepts in memory guide attention by deploying these templates, a phenomenon that we introduce as *learning-guided attention.* This work provides novel insights into the dynamic interplay between learning, memory, and attention.

**Significance Statement:** Extensive work shows that attention selects the most relevant information while learning new knowledge. Theoretically, the interaction between learning and attention is bidirectional; learned knowledge, in turn, guides attention to relevant information for that context. However, the mechanism by which knowledge in memory directs attention has remained largely unexplored. By developing an experimental paradigm that bridges the learning and attention literatures, we demonstrate that learning builds “attentional templates” which capture what is relevant in a specific learning context. While utilizing learned knowledge, individuals deploy these templates to allocate their attention to the most relevant information for a given situation. We introduce this phenomenon as *learning-guided attention*, providing novel insights into the dynamic interplay between learning, memory, and attention.

## 1. Introduction

*Concept learning* is the process of acquiring information that integrates multiple cognitive mechanisms; what is learned becomes *memory* and how it is learned depends on *attention*. Concepts are highly organized information structures that influence decision making, guide predictions about new experiences, and facilitate further learning (Seger et al., 2015; Zeithamova et al., 2012; Zeithamova & Bowman, 2020). Because of their critical role in shaping our understanding of the world, each concept is built with the information that is diagnostic for a particular context. This is an integral contribution of attention in learning: optimizing information sampling and ensuring successful concept formation by selecting what is relevant.

For many decades, attention has been formalized in prominent theories of concept learning (Kruschke, 1992; Love et al., 2004; Medin & Schaffer, 1978; Nosofsky, 1986; Shepard et al., 1961). Central to these theories is the notion that individuals learn to optimally allocate their attention to relevant features of a visual stimulus (Nosofsky, 1984). However, an unresolved but theoretically motivated question is that once concepts are built through the dynamic learning process, does optimized attention influence subsequent behaviour? We propose that concepts stored in memory guide our attention as a consequence of the learning experience in which tuned attention is represented in a context-specific manner. To evaluate this hypothesis, we leveraged an established phenomenon in the attention literature: attentional templates. These templates are specific to task demands that hold relevant spatial representations and provide top-down biases in perceptual processing and decision making (Bundesen, 1990). Here, we demonstrate that concept learning builds behaviourally relevant attentional templates that highlight the relevant features for a particular concept. Once learned, these concept-specific attentional templates can be flexibly deployed to guide our attention to relevant information.

Despite different perspectives on how concepts might be stored in memory, for instance, as prototypes (Smith & Minda, 1998), exemplars (Medin & Schaffer, 1978), or rules plus exceptions (Nosofsky et al., 1994; Palmeri & Nosofsky, 1995), attention is a fundamental component of many concept learning models. Attention is often evaluated with respect to its influence on concept-related decision making, maximizing the distinctions among the items of different concepts and minimizing differences among the items of the same concept (Nosofsky, 1984, 1986). According to the optimal attention allocation hypothesis, attention is equally distributed across all features of a visual stimulus early in learning, but it is gradually optimized to relevant features to minimize errors in concept judgments (Nosofsky, 1984) (Figure 1a).

**Figure 1.**
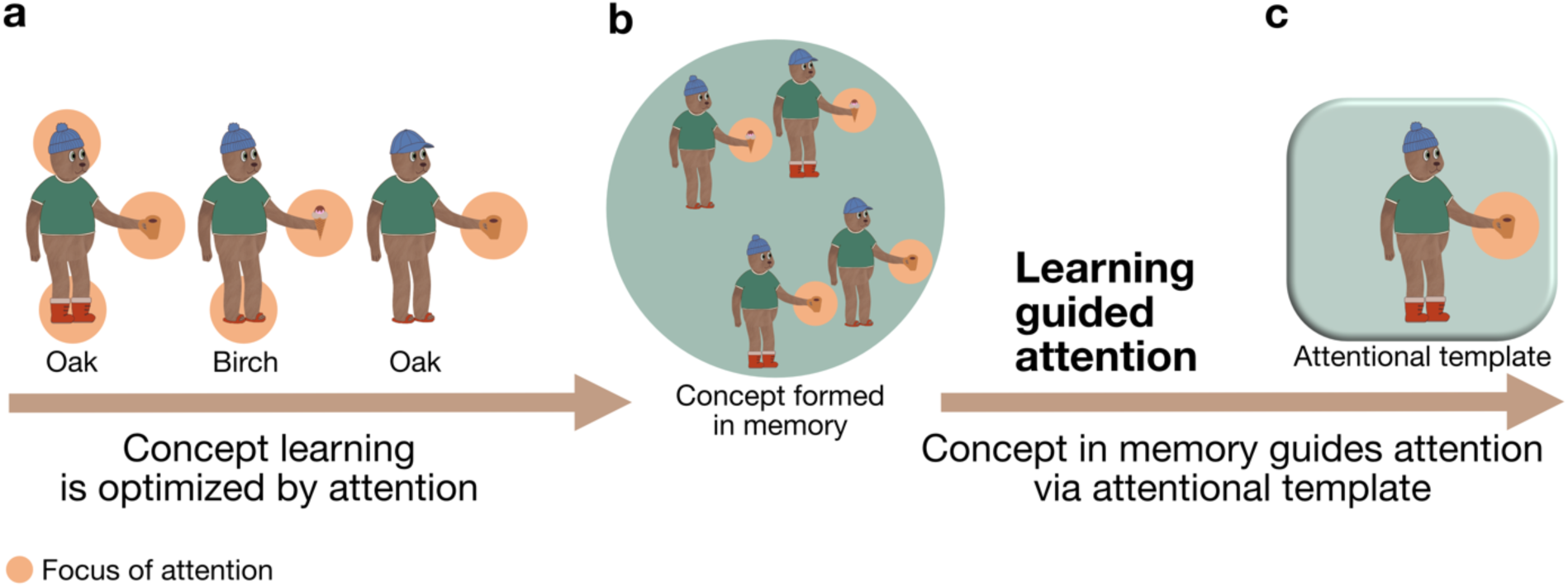
Learned concepts in memory guide attention by deploying corresponding attentional templates. a, Learning is optimized by attention. Attention during learning is tuned to organize visual input (e.g., bears belonging to the Oak versus Birch family) based on the features that separate categories (e.g., the food that the bears are holding). **b,** Successfully learned concepts are represented according to their relevant features (e.g., food) in memory, reflecting a shift in psychological space. **c,** We propose that concept learning builds attentional templates that represent prioritized information for a concept. This *learning-guided attention*, when invoked, will impact subsequent information sampling and behaviour.

Evidence of exogenous attention from eye-tracking supports this theory that attention is optimized in concept learning (Blair, Watson, & Meier, 2009; Blair, Watson, Walshe, et al., 2009; Rehder & Hoffman, 2005). This attentional tuning organizes the acquired information into concepts according to behaviourally relevant feature dimensions.

Many models of learning assume that concept members are represented in multidimensional psychological space based on their dissimilarity along feature dimensions (Palmeri & Gauthier, 2004). Before learning, items are represented in psychological space as a function of their shared features (Shepard et al., 1961). During learning, attentional weights for feature dimensions change relative to the learning context such that psychological space expands along the relevant feature dimensions and shrinks along the irrelevant dimensions (Kruschke, 1992; Nosofsky, 1984) (Figure 1b). This dynamic learning process influences our perception – the updated psychological space enhances the ability to perceive objects according to their respective concepts that extends beyond their simple perceptual similarities (Goldstone, 1994). Indeed, neuroimaging work provides empirical evidence that attention modulates stimulus representations in learning according to task demands, shaping how concepts are represented in memory (Mack et al., 2013, 2016, 2020). This close interaction between attention and memory is gaining theoretical prominence (Turner & Sloutsky, 2024). Recent models have conceptualized attention as a moment-by-moment information sampling process, wherein new information is constantly compared with previous experience in memory to both build concept representations and tune attention towards feature information that holds the most predicted utility for subsequent decisions (Braunlich & Love, 2022; Weichart et al., 2022). Despite extensive work on how attention influences the learning process and resulting memory representations, the impact of optimized attention on subsequent behaviour following learning remains unexplored.

Foundational theories of attention posit that attention is guided by mental representations of the task context that are stored in *attentional templates* (Bundesen, 1990; Desimone & Duncan, 1995). These templates contain goal-relevant information about features (e.g., size, shape, and colour of a target) which prioritize spatial locations in the visual field. Such top-down attentional processes (Bacon & Egeth, 1994) along with bottom-up saliency (Theeuwes, 1991, 1992) are the two major factors in attention guidance. However, beyond this classical dichotomy of attention, the sampling of experience over time, namely selection history, can also influence attention (Anderson et al., 2021; Awh et al., 2012). Priming (Maljkovic & Nakayama, 1994), contextual cueing (Chun & Jiang, 1998), and statistical learning (Jiang et al., 2013) are examples of selection history that cannot be explained solely by current goals or physical saliency. Selection history rather reflects how attention was allocated in the past and prioritizes the current context that best matches the previous experience. This is well captured in location probability cueing; eye movements are faster to the location where the target had a higher probability of appearing in statistical learning (Jiang et al., 2014). Nevertheless, selection history, even in an implicit setting, is intertwined with top-down attentional goals such that it builds attentional templates over experiences and instates them when necessary (Zhang & Carlisle, 2023). Although attentional templates are traditionally considered to be maintained in working memory (Bundesen, 1990; Bundesen et al., 2005; Desimone & Duncan, 1995), accumulating evidence suggests their eventual transfer to long-term memory (Carlisle et al., 2011; Woodman et al., 2007; Zhang & Carlisle, 2023). This raises an intriguing question of whether concept learning also builds behaviourally relevant attentional templates across learning experiences. Such templates could later be deployed to guide attention as we retrieve information regarding a particular concept.

We propose that concept learning results in more than the organization of concept representations in memory, but also the formation of concept-specific attentional templates that represent prioritized feature information. As concepts are stored in memory, so are the attentional templates. Critically, these attentional templates, once learned, can be flexibly deployed to efficiently guide attention to relevant features. To test this proposal, participants first learned to categorize the same visual stimuli according to two different category structures before being tested with a novel probe task. To briefly foreshadow, after learning we observe a distinct benefit in processing unrelated visual targets appearing in concept-relevant feature locations, a hallmark of attentional guidance (Posner, 1980). This *learning-guided attention* effect provides a unique window in the dynamic interplay between learning, memory, and attention.

## 2. Methods

### 2.1. Participants

This study included 194 participants (116 females) with the mean age of 18.83 ± 1.22. The study was approved by the University of Toronto Research Ethics Board. Participants were recruited from an introductory psychology course at the University of Toronto. All participants had normal or corrected-to-normal vision and normal colour vision. Written informed consent was collected from all participants prior to their participation. Participants received course credit for their participation.

### 2.2. Study Stimuli

We designed a novel stimulus set of eight cartoon bears. The base bear image was drawn in a standing position with head, body, extremities and a green t-shirt (Figure 2a). The base bear image was consistent across the eight stimuli. Three features of the bear varied across the learning tasks, specifically, the hat it was wearing, the food it was holding, and the shoes it had on (Figure 2a). Each feature dimension had two values; the hat could be a tuque or cap, the food could be ice cream or a coffee cup, and the shoes could be boots or sandals (Figure 2a). Notably, the bear stimulus set was designed to include spatial and feature-based properties. Each feature dimension included similar colour content: hats were blue, food was mostly brown, and shoes were red. Additionally, these bear features (i.e., hat, food, shoes) were at an equal distance from the centre of the bear. The combination of the three features, each with binary values, yielded eight different bear images. Bear images were 1200 x 1200 pixels in size and presented on a white background at the centre of the screen.

**Figure 2.**
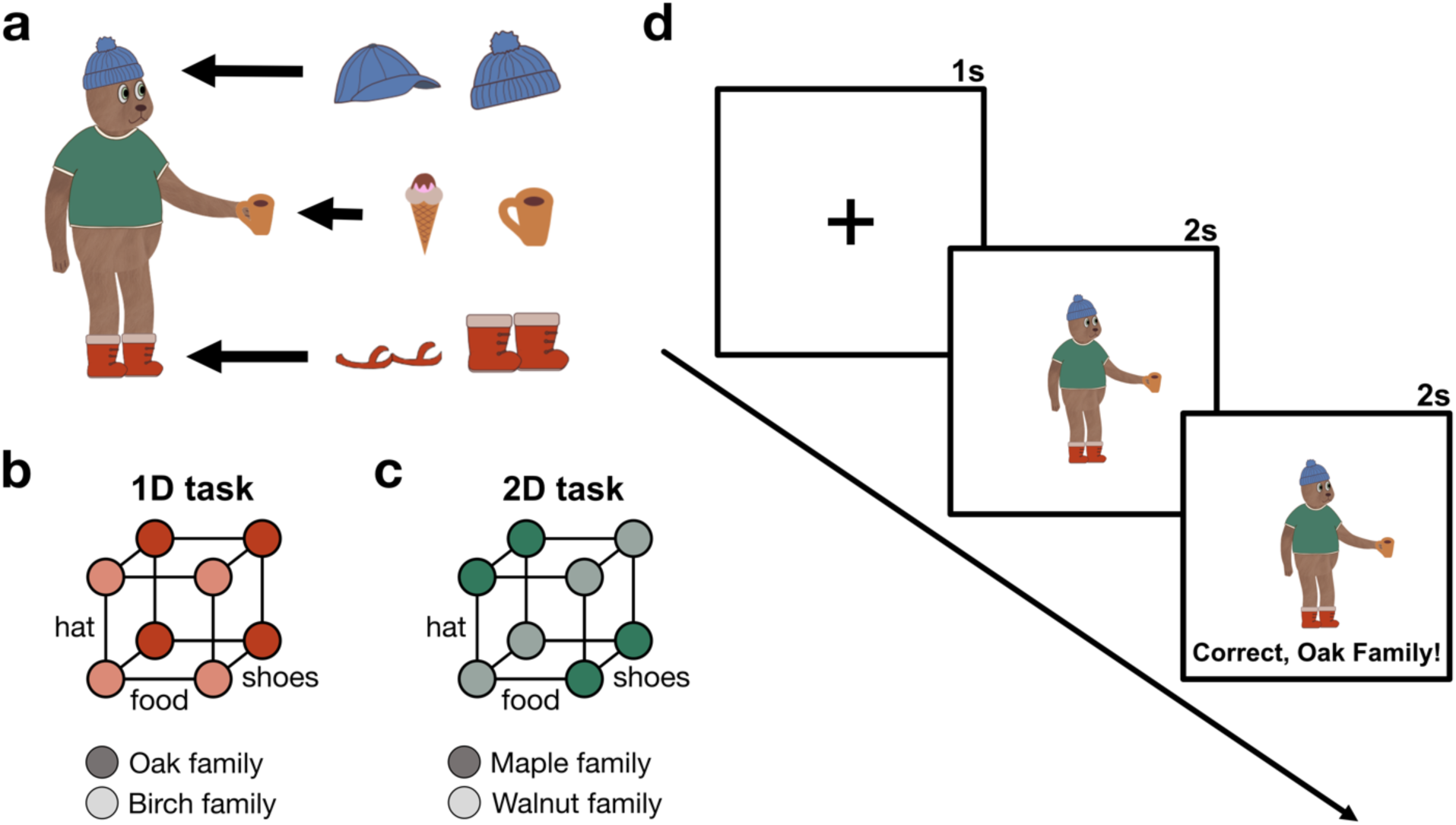
Stimulus set and study design in concept learning tasks. a, The base bear image included 3 varying features: the hat could be a cap or a tuque, the food could be an ice cream or a coffee cup, and the shoes could be sandals or boots. **b,** The one-dimensional task (1D task) rule was based on one feature dimension that separated the bears into Oak and Birch families. **c,** The two-dimensional (2D task) task included two separate bear families, namely Maple and Walnut, that were based on the combination of two feature dimensions. **d,** Learning trials were identical in both learning tasks but included different concept rules. Each learning trial started with a 1s-long fixation cross, that was followed by the bear stimulus for 2s, during which participants were instructed to decide the family of the bear. Participants immediately received feedback, where a verbal text as well as the bear stimulus were presented for 2s.

### 2.3. Study Procedure

The study included three blocks: two separate category learning tasks and a final test phase with mixed probe trials. Participants were instructed to categorize the bear stimuli into the families they belonged to according to the 3 varying features (i.e., hat, food, shoes). The two learning tasks utilized the same bear images, but the categories for the two tasks were defined by different feature dimensions. Participants were not told what the concept rules were or the features that mattered for a given task. Instead, they learned to categorize the bears based on trial-by-trial feedback. Following the learning blocks, participants completed a test phase in which they categorized the bears similar to the learning tasks but did not receive any feedback. The test block included probe trials that were leveraged to assess attention. They were asked to respond as quickly as possible across all three blocks of the experiment. The experiment was designed and coded in PsychoPy2, version 2022.2.5 (Peirce et al., 2019).

### 2.4. Concept Learning Tasks

One of the learning tasks included a one-dimensional rule (i.e., 1D task) in which the binary values of one feature determined the bear categories, for instance a coffee cup versus ice cream (Figure 2b). Participants were told that each bear belonged to either the Oak or Birch family.

They used the keyboard to respond and received feedback on each trial such as “Correct, Oak family!” or “Incorrect, Birch family!”. The other learning task included a two-dimensional rule (i.e., 2D task), specifically exclusive-or (XOR), categories such that the separation between the bear categories was based on the values of 2 different features (Figure 2c). In this task, the bears belonged to Maple or Walnut families. Similar to the 1D task, participants received feedback on each trial. Each learning task included 80 trials with 10 repetitions of each bear. The bear stimulus was displayed for 2s on the screen, during which participants were instructed to respond (Figure 2d). They received feedback immediately; the text feedback appeared right below the bear stimulus, indicating whether the response was correct and incorrect (Figure 2d). The feedback was presented for 2s. The interstimulus interval was 1s, during which a fixation cross appeared at the centre of the screen.

Concept rules in each learning task were defined based on different feature dimensions. For instance, if the hat was the determining feature for the 1D task, then the combination of the food and shoes determined the 2D task. This design separated the attentional demands of the two concept learning tasks. Which feature dimensions were mapped to the different concept rules was randomized across participants. The order of the learning tasks was also randomized across participants. Before starting the second learning task, participants were reminded that the same bears were going to appear on the screen, but, this time, they needed to categorize the bears differently as there were two new families in this study block.

### 2.5. Test Phase with Probe Trials

The test block included two types of trials: test and probe trials. Probe trials were implemented as an independent task to capture attentional templates (Figure 3a). There were 248 trials in total with 128 test and 120 probe trials. Each trial started with one of the two possible cues, indicating which category task should be performed on that trial: “Oak or Birch?” or “Maple or Walnut?” (Figure 3a). The cue was presented for 2s followed by a fixation cross for 0.5s. On test trials, the bear stimulus was then displayed for 3s. Participants were instructed to categorize the bears according to the cued concept learning task. No feedback was provided on test trials. Of the 128 test trials, 64 of them were related to the first learning task, and the other 64 were about the second learning task. Each bear stimulus was tested 8 times, and all test trials were randomized within a participant.

**Figure 3.**
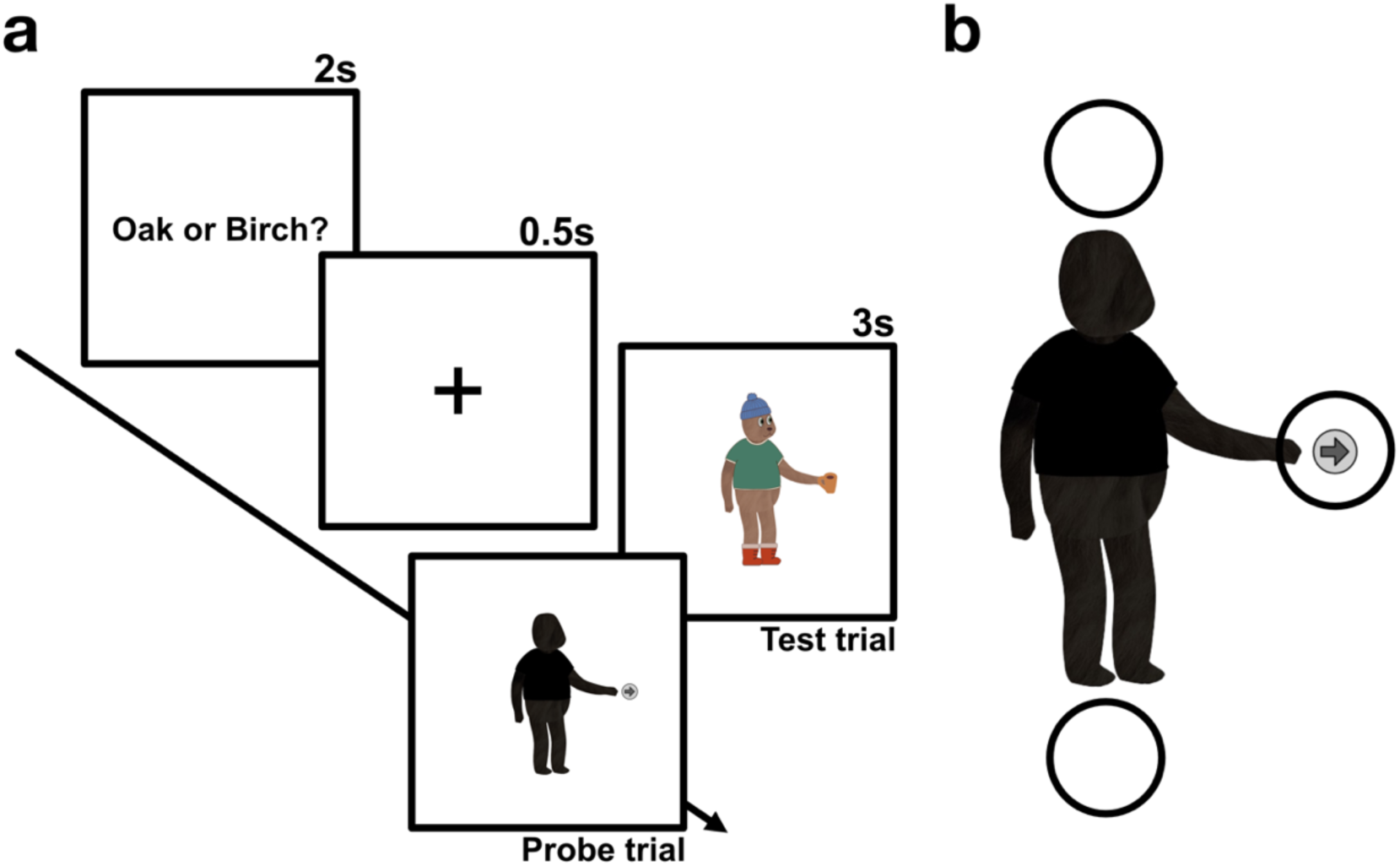
Test block with probe trials as an independent task to assess attentional templates. a, Participants were cued to one of the learning tasks on each trial in the test phase for 2s. Then, a 0.5s fixation cross was followed by either a test or probe trial. On test trials, participants were instructed to categorize the bear according to the cued learning task but did not receive feedback. On probe trials, they instead encountered a shadow bear, where they had to respond if the arrow pointed right or left. **b,** Probe trials did not include any bear features, instead a right or left arrow appeared in one of the three feature locations. The circles represent arrow locations but were not present in the experiment. Valid probes included an arrow in the feature location that was relevant to the cued learning task. The arrow on invalid probes was at an irrelevant feature location with respect to the cued learning task on that trial.

On a random subset of trials, a probe appeared instead of the bear stimulus following the cue (Figure 3b). Probes were composed of a shadow bear with the same shape and size as the bear stimulus that was used in the learning tasks but did not include any of the visual features or colours (Figure 3b). Instead, a small arrow appeared in one of the feature locations: next to the head, hand, or feet of the bear (Figure 3b). All arrow locations had the same distance to the centre of the bear. On 50% of the probe trials, an arrow appeared at a feature location that was relevant to the cued concept learning task. We call these “valid” probe trials. The remaining 50% of the trials included “invalid” probes such that an arrow appeared at a location that was irrelevant to the cued category learning task. The arrow pointed either right or left, with an equal number of right and left arrows appearing at each feature location across the probe trials.

Participants were instructed to respond whether the arrow was pointing right or left when they encountered these shadow bears. There was no feedback on probes.

### 2.6. Assessment of Learning Performance

All analyses were performed with R statistical software, version 4.2.1 (R Core Team, 2021). Participants’ learning curves across 80 trials were examined by calculating their percent correct on each trial in each of the learning tasks. Their learning accuracy was calculated based only on the second half of the learning tasks: the number of correctly categorized bears divided by 40 trials. These learning accuracies were used to separate participants into learner versus non-learner groups in each concept learning task. The learning threshold was calculated based on a random guessing model; with 80 trials and 5000 permutations, the probability of performing above the random guessing model with a 95% confidence interval was 61.25%. Leveraging this threshold, participants were labeled as learners if they reached an accuracy of 61.25% or above. If they performed below this learning threshold, they were grouped into non-learners. The same learning threshold was separately applied to the two learning tasks.

### 2.7. Assessment of Concept Specific Attentional Templates

Deployment of attentional templates was evaluated based on the response time to the valid versus invalid probes in the test phase. Participants who responded to 15 or more probes in less than 250ms were excluded, and thus 12 participants were excluded from the attentional template-specific analyses (N=182). We also excluded the probe trials where the response time was under 150ms. We used a generalized linear mixed effects model with an inverse gaussian family distribution and identity link function (Lo & Andrews, 2015) to predict trial-by-trial response times for probes with an interaction between the feature location (i.e., relevant/irrelevant), learner type (i.e., learner/non-learner), and task type (i.e., 1D/2D task) with random effects for participants and trials (*lme4* version 1.1-30, *lmerTest* version 3.1-3) (Bates et al., 2015; Kuznetsova et al., 2017). Only correct responses to probes were included in these estimates. To investigate whether attentional guidance varies with the complexity of the learning task, we set up a separate generalized linear mixed effects model but included only the learners. This enabled the comparison of the response times to the valid probes for the 1D versus 2D task.

### 2.8. Relating Attentional Template Deployment to Learning Performance

Beyond subgroup analyses (i.e., learners versus non-learners), we assessed the link between learning performance and resulting attention guidance on a continuum. We computed each participant’s median response time to valid and invalid probes for both learning tasks. These calculations were based only on the correct probe trials. We calculated the response time benefit for each participant by subtracting their median response time for valid probes from that of invalid probes for each learning task. The distribution difference between the 1D and 2D tasks was assessed with a paired t-test. We expected that the response time benefit would be greater with successful learning, indicating that efficient deployment of attentional templates relies on the learning experience. We tested this hypothesis by setting up a linear mixed effects model, where the response time benefit of each participant was predicted by their learning accuracy in each learning task. We examined the correlation between response time benefit and learning accuracy based on the model predictions.

## 3. Results

### 3.1. Individual Variability in Concept Learning

The 1D task was quickly learned, but the performance in the 2D task was relatively lower, as observed in the learning curves (Figure 4a). Overall, participants learned the 1D task very well (M = 89.32%, SD = ±18.04; Figure 4b). Approximately 87% of the participants performed at or above the learning threshold and were labeled as learners in the 1D task. In the 2D task, learning performance was lower but highly variable (M = 63.50%, SD = ±21.87; Figure 4b). About 47% of the participants reached the learning threshold and were labeled as learners in the 2D task.

**Figure 4.**
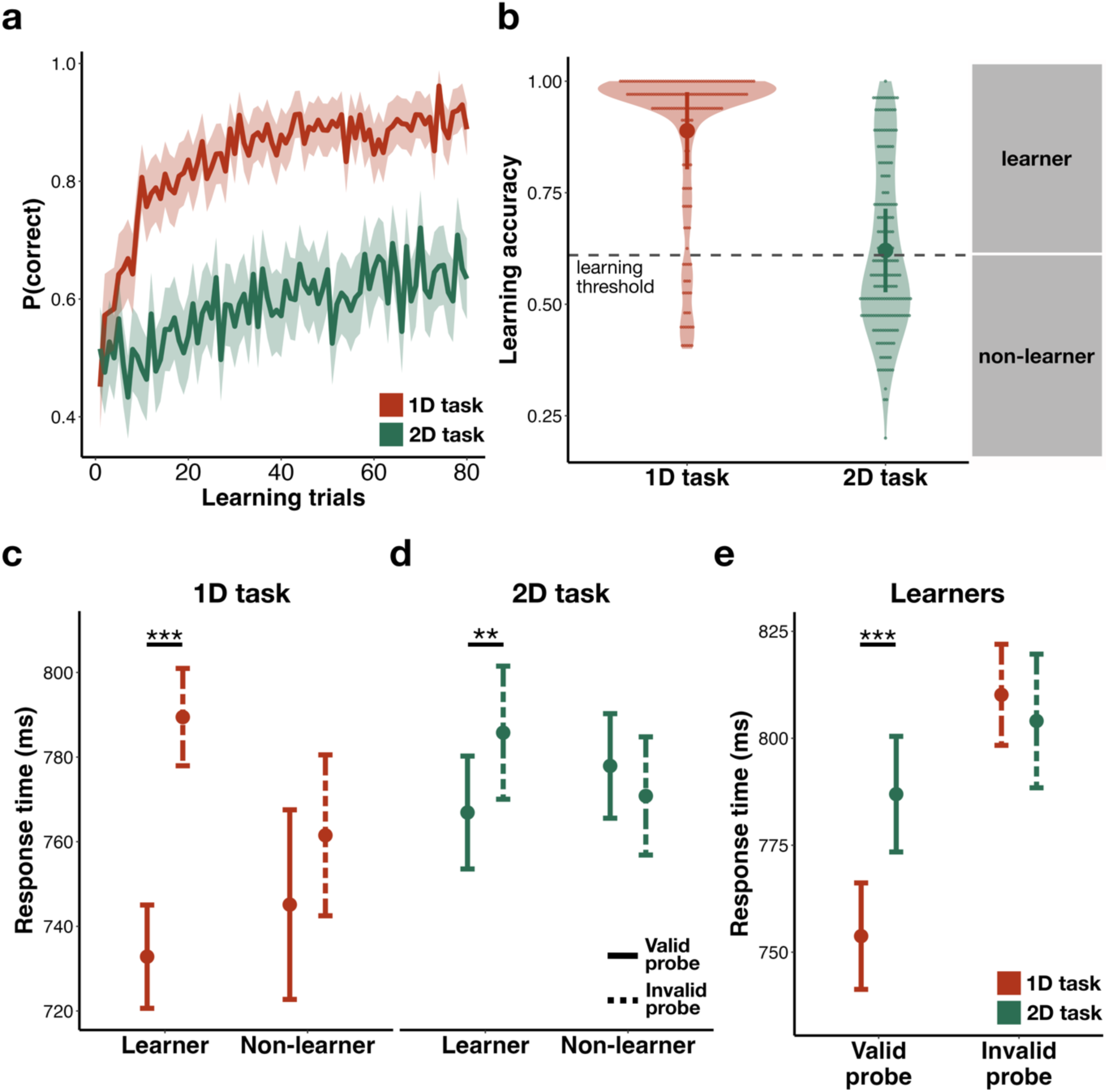
Deployment of attentional templates that emerged from successful learning was captured with probes. a, Average probability of correct responses across learning trials in both the 1D and 2D tasks. The ribbon represents the 95% confidence interval of the mean. **b,** Participants’ learning performance was calculated based on the second half of the learning trials. Each dot is an individual learning accuracy. Error bars present the mean and standard deviation. The 2D task exhibited relatively greater learning variability. Participants were separated into learners (at or above 61.25%) and non-learners (below 61.25%) based on the learning threshold (gray dotted horizontal line). **c,** Model predictions were plotted with the mean (the centre) as well as the error bars of 95% of confidence interval. Learners were significantly faster to respond to valid (i.e., dashed) probes than invalid (i.e., dotted) probes in both the 1D and **d,** 2D tasks. The response time differences between valid and invalid probes were not observed in non-learners. **e,** Learners responded significantly faster to valid probes in the 1D than the 2D task. *Note. ** p<.01, *** p<.001*

### 3.2. Concept-Specific Attentional Templates Captured with Response Time

We aimed to identify the differences between the learners and non-learners in both task with respect to their response times to probe trials as a measure of attentional template deployment. Overall accuracy on probe trials was high in both the 1D (M = 88.70%, SD = ±14.28) and 2D tasks (M = 87.90%, SD = ±14.50). These probe trials were designed as an independent task from the learning tasks as participants responded to the arrow rather than categorizing the bears. The main hypothesis was that the concept-specific attentional template that is activated by the cue would have a carry-over attention effect on the shadow bear, allowing us to assess attentional bias in an independent setting. We expected that if the arrow appeared in the feature location that was relevant for the cued task (i.e., valid probes), the response time would be faster to the probes, indicating the deployment of attentional templates, than if the arrow was at an irrelevant location (i.e., invalid probes). Confirming our hypothesis, when participants were cued to the 1D task in the test phase, learners were significantly faster to respond to the valid probe trials than invalid ones (β =-56.62, SE = 4.62, z =-12.25, p <.001; Fig. 4c). Non-learners did not show a response time difference for valid versus invalid probes (β =-16.38, SE = 11.33, z =-1.45, p = 0.15; Fig. 4c), indicating that no attentional guidance emerged from concept learning. Similarly, when participants were cued to the 2D task, learners responded significantly faster to the valid probes than invalid ones (β =-18.87, SE = 7.01, z =-2.69, p <.01; Fig. 4d). Again, this response time difference was not observable in the non-learners for the 2D task (β = 7.10, SE = 5.93, z = 1.20, p = 0.23; Fig. 4d). Additionally, we observed that the learners were significantly faster to respond to valid probes for the 1D task than those for the 2D task (β =-33.19, SE = 5.58, z = - 5.95, p <.001; Fig. 4e). This supports our hypothesis that attentional guidance varies with the complexity of the learning task.

### 3.3. Response Time Benefit of Attentional Templates Linked to Successful Learning

The response time benefit was greater on probes in the 1D task (M = 59.87, SD = ±84.85) than in the 2D task (M =-0.015, SD = ±78.10), t (181) = 6.86, p<.0001 (Figure 5a). Learning accuracy in both the 1D (β=76.22, 95% CI [14.33, 138.10], p <.05) and 2D tasks (β=70.74, 95% CI [13.53, 127.94], p <.05) was significantly associated with the response time benefit measured on probe trials, highlighting that learning success was linked to the efficiency and flexibility of the attentional guidance observed on probe trials (Figure 5b).

**Figure 5.**
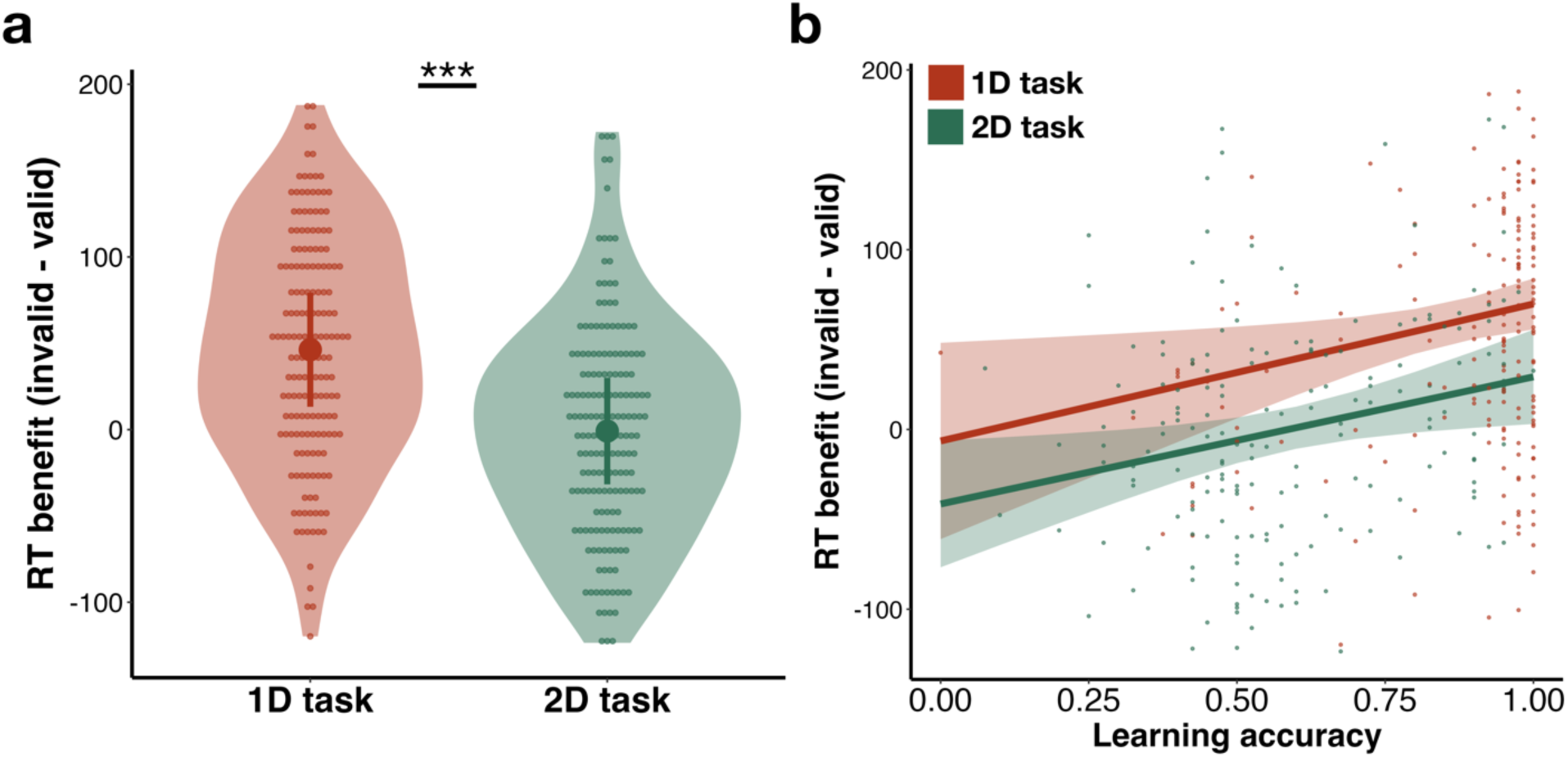
Efficient deployment of attentional templates relates to learning success. **a,** Response time benefit was calculated for each participant in each learning task by subtracting their median response time for valid probes from that of invalid probes. Dots represent the individual response time benefits. Error bars are the 95% confidence interval plotted around the mean. The 1D task showed significantly greater response time benefit than the 2D task. **b,** Individual learning accuracies were related to the response time benefit measured on probe trials, indicating that efficient deployment of attentional templates relied on learning success. Model predictions of the relationship are depicted with solid lines and ribbons, representing 95% confidence intervals. Dots represent individual participant data. *Note. *** p<.001*

## 4. Discussion

We present compelling evidence for the emergence of behaviourally relevant attentional templates in concept learning. Established theories and computational models of concept learning emphasize the optimization of attention to relevant features to form a particular concept.

Our findings reveal that this dynamic interplay between learning and attention extends significantly beyond concept formation. We demonstrate that concepts, once stored in memory, actively guide subsequent attentional deployment. Through measurements of processing speed to probes following learning, particularly within task-switching contexts, we captured the operation of these concept-specific attentional templates. Notably, learners who successfully acquired concepts responded faster to valid probes at task-relevant feature locations compared to invalid probes that were at irrelevant feature locations, signifying the deployment of these specialized templates. This underscores that learning constructs attentional templates by prioritizing the very feature information critical for a given concept. Because such templates are integrated into conceptual knowledge, they exert a lasting influence on behaviour. Furthermore, we observed that the efficacy of learning dictates the efficiency of attentional template deployment: greater attention optimization during learning translated into a more substantial response time benefit during concept retrieval. Consequently, the degree to which learned concepts direct our attention is intrinsically linked to the dynamic learning process through which attentional templates are forged to encapsulate concept-relevant representations. We introduce this novel phenomenon as *learning-guided attention*.

The notion of attentional templates is well-established in perceptual tasks such as visual search (Conci et al., 2019; van Moorselaar et al., 2014; Wolfe, 2021; Yu & Geng, 2019). The nature of these templates is determined not only by information actively maintained in working memory (e.g., the current search target) but also by associated information held in long-term memory.

Such long-term memory contributions can arise from the environmental context, as demonstrated in the extensive contextual cueing literature wherein repeated spatial configurations implicitly guide attention (Jiang et al., 2019) or form more explicit top-down strategic goals (Zhang & Carlisle, 2023). Our research significantly extends these findings to concept learning by demonstrating that explicit learning goals, instantiated through the process of concept acquisition, profoundly shape the characteristics of attentional templates. This observation resonates with, yet distinctively expands upon, the classical contextual cueing paradigm, where prioritized spatial information, developed through repeated experiences, accelerates response times (Jiang et al., 2019; Zhang & Carlisle, 2023). Contextual cueing is an implicit mechanism; repeated exposure to a search context allows for the extraction of regularities, such as target location, which then facilitates attention deployment. While both phenomena demonstrate attention deployment related to selection bias, *learning-guided attention* in our study arises from concept formation processes, in which learning instances are organized into knowledge structures. The attentional guidance here is not solely reliant on search history or spatial configurations. Instead, it involves the integration of multidimensional information, encompassing diagnostic features, their characteristics, and category labels, which together structure concepts and support generalization to new instances. Thus, encountering information that matches an existing, explicitly learned concept guides attention to its relevant features precisely because attention was optimized for these feature dimensions during initial learning.

This framework details the mechanism by which concept-specific templates direct attentional biases during retrieval. Thus, many episodic experiences during concept learning contribute to developing behaviourally relevant attentional templates, where attention optimized during learning exerts a protracted influence on subsequent behaviour, even after learning is complete.

Attentional guidance by learned concepts operates as a more explicit and complex mechanism than implicit processes like contextual cueing, despite superficial similarities. Central to effective concept learning, attention enables the flexible selection of feature information tailored to varying learning goals (Nosofsky, 1984, 1986). We therefore hypothesized that if concept learning utilizes flexible attentional templates to prioritize distinct information for different objectives, then changing the concept decision task would trigger specific, corresponding shifts in attention. Our results confirmed this, demonstrating that such flexible attentional templates guide attention differently across diverse learning tasks, even when stimuli remain constant. A key piece of evidence supporting this is the flexible deployment of attentional templates contingent on the complexity of the learning task and the specific conceptual distinctions required. We observed that when participants learned to categorize the same stimuli according to different rules (e.g., a 1D vs. 2D rule), the resulting attentional templates were specifically tailored to the demands of each task. Attention is a critical mechanism for successful concept learning, enabling the flexible selection of feature information pertinent to different learning demands. This was further evidenced by the magnitude of the response time benefit, which scaled with the degree of attentional focus required: a larger benefit was observed for tasks demanding attention to a single feature dimension (i.e., 1D) compared to tasks requiring more diffuse attention across multiple dimensions (i.e., 2D). This suggests that the precision of the attentional template, and thus its benefit, is modulated by the attentional demands inherent in the learning task. If concept learning is indeed driven by such flexible attentional templates that guide the selection of distinct information for varying learning goals, then shifting the conceptual decision should, and did, result in concept-specific impacts on attentional deployment.

Attention is often characterized as a modulatory factor in the learning process, whereby attentional weights assigned to feature dimensions are adjusted via feedback (Kruschke, 1992; Love et al., 2004; Nosofsky, 1986). Our results suggest that attentional templates originating from concept learning embody this reconfigured psychological space, reflecting the dimensional stretching along diagnostic features that occurs during concept formation (Gauthier & Palmeri, 2002). Specifically, we showed that successfully learning to categorize identical stimuli in multiple ways necessitates an effective orthogonalization of psychological space, where attentional templates, by prioritizing the most diagnostic information for a given goal, play a crucial role in this representational differentiation. This aligns with contemporary views of attentional templates as customized, predictive representations, not mere replicas of a target, but rather entities shaped by expectations and learned perceptual biases (Geng & Witkowski, 2019). We propose that the templates emerging from concept learning share these characteristics. They develop throughout the learning process, capturing tuned attentional weights and the modified concept space, thereby enabling learned concepts in memory to guide attention effectively during retrieval, highlighting the reliance of this guidance on the dynamic learning experience itself.

The present findings are in accord with, but also significantly advance, the broader field of memory-guided attention, which studies the influence of past experiences on current attentional deployment (Chen & Hutchinson, 2019; Hutchinson & Turk-Browne, 2012; Moores et al., 2003). Memory-guided attention encompasses phenomena in which, for instance, prior encounters with an object at a specific location, or an object’s learned association with reward, can bias attention towards that object or location. While both memory-guided attention and our learning-guided attention highlight the influence of long-term memory on attention, distinct from purely top-down strategic control or bottom-up stimulus salience, there are critical differences. Memory-guided attention often refers to biases arising from specific episodic memories, priming, or semantic memories (Chen & Hutchinson, 2019; Hutchinson & Turk-Browne, 2012). Learning-guided attention, on the other hand, emerges from the dynamic tuning of attention and the concurrent modification of memory representations during new concept formation. The attentional templates related to learning-guided attention are not just linked to a prior location or a specific past event; they are tuned to the diagnostic features that define the concept itself, features that were actively weighted and prioritized during the learning process to enable categorization and generalization. Furthermore, this concept-specific attention optimization in learning generates distinct attentional templates for the exact same input based on task demands, unlike memory-guided attention in which the source of attentional bias is often less apparent (Chen & Hutchinson, 2019; Hutchinson & Turk-Browne, 2012). Thus, while memory-guided attention demonstrates that past experience can guide attention, learning-guided attention specifies a mechanism in which the act of learning a concept systematically builds a feature-based attentional template that reflects the conceptual structure. The templates in learning-guided attention are a direct consequence of the attentional operations necessary for successful concept acquisition.

A thorough examination of the intricate relationship between learning, memory, and attention calls for a multidisciplinary perspective, as adopted in our work. While classical theories have explored how attention refines concept learning and shapes the resulting representations, the influence of learned concepts on subsequent attentional deployment has remained comparatively unexplored. We argue that attention, once tuned to relevant information during concept formation, has enduring effects on behaviour. Specifically, the novel concept of *learning-guided attention* posits that concept-specific attentional templates, developed during learning, capture prioritized feature information. These templates, stored alongside the concepts in memory, are then deployed upon concept retrieval to allocate attention to relevant features. The distinct nature of these templates – explicit, feature-based, and conceptually-driven – sets them apart from traditional probability or contextual cueing paradigms. This distinction arises because they are actively shaped by learning goals, rather than resulting from the monotonous accumulation of statistical information. Importantly, our work integrates insights from concept learning theories regarding attention optimization, findings from memory-guided attention research on the influence of past experiences, and the classic attention literature on attentional templates.

Understanding *learning-guided attention* is, therefore, a critical step in comprehending the dynamic and reciprocal interplay between learning, memory, and attention.

## 5. Data Availability

Data and the stimulus set will be available upon request. All inquiries can be directed to the corresponding author, Melisa Gumus, at.

## 6. Acknowledgements

This project was funded by Natural Sciences and Engineering Research Council (NSERC) Discovery Grants (RGPIN-2017-06753 to MLM and RGPIN-2024-0588 to MLM), Canadian Institute of Health Research (CIHR) Grant (PJT-178337 to MLM), Brain Canada Foundation Grant (to MLM), and Vanier Canada Graduate Scholarship provided by NSERC (to MG). We also would like to thank Wen Jia (Michelle) Zhao and Julia Do for their help with data collection.

## 7. Author Contributions

M.G. and M.L.M. designed the study. M.G. and Z.Z.Y.L. collected the data. M.G. created the stimulus set, conducted the analyses, and wrote the manuscript. M.L.M. supervised the project and provided advice. All authors edited the manuscript.

## References

Anderson, B. A., Kim, H., Kim, A. J., Liao, M.-R., Mrkonja, L., Clement, A., & Grégoire, L. (2021). The past, present, and future of selection history. Neuroscience & Biobehavioral Reviews, 130, 326–350. 10.1016/j.neubiorev.2021.09.004

Awh, E., Belopolsky, A. V., & Theeuwes, J. (2012). Top-down versus bottom-up attentional control: A failed theoretical dichotomy. Trends in Cognitive Sciences, 16(8), 437–443. 10.1016/j.tics.2012.06.010

Bacon, W. F., & Egeth, H. E. (1994). Overriding stimulus-driven attentional capture. Perception & Psychophysics, 55(5), 485–496. 10.3758/BF03205306

Bates, D., Mächler, M., Bolker, B., & Walker, S. (2015). Fitting Linear Mixed-Effects Models Using lme4. Journal of Statistical Software, 67, 1–48. 10.18637/jss.v067.i01

Blair, M. R., Watson, M. R., & Meier, K. M. (2009). Errors, efficiency, and the interplay between attention and category learning. Cognition, 112(2), 330–336. 10.1016/j.cognition.2009.04.008

Blair, M. R., Watson, M. R., Walshe, R. C., & Maj, F. (2009). Extremely selective attention: Eye-tracking studies of the dynamic allocation of attention to stimulus features in categorization. *Journal of Experimental Psychology: Learning*, Memory, and Cognition, 35(5), 1196–1206. 10.1037/a0016272

Braunlich, K., & Love, B. C. (2022). Bidirectional influences of information sampling and concept learning. Psychological Review, 129(2), 213–234. 10.1037/rev0000287

Bundesen, C. (1990). A theory of visual attention. Psychological Review, 97(4), 523–547. 10.1037/0033-295x.97.4.523

Bundesen, C., Habekost, T., & Kyllingsbæk, S. (2005). A Neural Theory of Visual Attention: Bridging Cognition and Neurophysiology. Psychological Review, 112(2), 291–328. 10.1037/0033-295X.112.2.291

Carlisle, N. B., Arita, J. T., Pardo, D., & Woodman, G. F. (2011). Attentional Templates in Visual Working Memory. The Journal of Neuroscience, 31(25), 9315–9322. 10.1523/JNEUROSCI.1097-11.2011

Chen, D., & Hutchinson, J. B. (2019). What Is Memory-Guided Attention? How Past Experiences Shape Selective Visuospatial Attention in the Present. In T. Hodgson (Ed.), Processes of Visuospatial Attention and Working Memory (pp. 185–212). Springer International Publishing. 10.1007/7854_2018_76

Chun, M. M., & Jiang, Y. (1998). Contextual Cueing: Implicit Learning and Memory of Visual Context Guides Spatial Attention. Cognitive Psychology, 36(1), 28–71. 10.1006/cogp.1998.0681

Conci, M., Deichsel, C., Müller, H. J., & Töllner, T. (2019). Feature guidance by negative attentional templates depends on search difficulty. Visual Cognition, 27(3–4), 317–326. 10.1080/13506285.2019.1581316

Desimone, R., & Duncan, J. (1995). Neural mechanisms of selective visual attention. Annual Review of Neuroscience, 18, 193–222. 10.1146/annurev.ne.18.030195.001205

Gauthier, I., & Palmeri, T. J. (2002). Visual neurons: Categorization-based selectivity. Current Biology: CB, 12(8), R282–284. 10.1016/s0960-9822(02)00801-1

Geng, J. J., & Witkowski, P. (2019). Template-to-distractor distinctiveness regulates visual search efficiency. Current Opinion in Psychology, 29, 119–125. 10.1016/j.copsyc.2019.01.003

Goldstone, R. (1994). Influences of categorization on perceptual discrimination. Journal of Experimental Psychology. General, 123(2), 178–200. 10.1037//0096-3445.123.2.178

Hutchinson, J. B., & Turk-Browne, N. B. (2012). Memory-guided attention: Control from multiple memory systems. Trends in Cognitive Sciences, 16(12), 576–579. 10.1016/j.tics.2012.10.003

Jiang, Y. V., Sisk, C. A., & Toh, Y. N. (2019). Implicit guidance of attention in contextual cueing: Neuropsychological and developmental evidence. Neuroscience & Biobehavioral Reviews, 105, 115–125. 10.1016/j.neubiorev.2019.07.002

Jiang, Y. V., Swallow, K. M., Rosenbaum, G. M., & Herzig, C. (2013). Rapid acquisition but slow extinction of an attentional bias in space. Journal of Experimental Psychology. Human Perception and Performance, 39(1), 87–99. 10.1037/a0027611

Jiang, Y. V., Won, B.-Y., & Swallow, K. M. (2014). First saccadic eye movement reveals persistent attentional guidance by implicit learning. Journal of Experimental Psychology. Human Perception and Performance, 40(3), 1161–1173. 10.1037/a0035961

Kruschke, J. K. (1992). ALCOVE: An exemplar-based connectionist model of category learning. Psychological Review, 99(1), 22–44. 10.1037/0033-295X.99.1.22

Kuznetsova, A., Brockhoff, P. B., & Christensen, R. H. B. (2017). lmerTest Package: Tests in Linear Mixed Effects Models. Journal of Statistical Software, 82, 1–26. 10.18637/jss.v082.i13

Lo, S., & Andrews, S. (2015). To transform or not to transform: Using generalized linear mixed models to analyse reaction time data. Frontiers in Psychology, 6. 10.3389/fpsyg.2015.01171

Love, B. C., Medin, D. L., & Gureckis, T. M. (2004). SUSTAIN: A network model of category learning. Psychological Review, 111(2), 309–332. 10.1037/0033-295X.111.2.309

Mack, M. L., Love, B. C., & Preston, A. R. (2016). Dynamic updating of hippocampal object representations reflects new conceptual knowledge. Proceedings of the National Academy of Sciences, 113(46), 13203–13208. 10.1073/pnas.1614048113

Mack, M. L., Preston, A. R., & Love, B. C. (2013). Decoding the brain’s algorithm for categorization from its neural implementation. Current Biology: CB, 23(20), 2023–2027. 10.1016/j.cub.2013.08.035

Mack, M. L., Preston, A. R., & Love, B. C. (2020). Ventromedial prefrontal cortex compression during concept learning. Nature Communications, 11(1), Article 1. 10.1038/s41467-019-13930-8

Maljkovic, V., & Nakayama, K. (1994). Priming of pop-out: I. Role of features. Memory & Cognition, 22(6), 657–672. 10.3758/BF03209251

Medin, D. L., & Schaffer, M. M. (1978). Context theory of classification learning. Psychological Review, 85(3), 207.

Moores, E., Laiti, L., & Chelazzi, L. (2003). Associative knowledge controls deployment of visual selective attention. Nature Neuroscience, 6(2), 182–189. 10.1038/nn996

Nosofsky, R. M. (1984). Choice, similarity, and the context theory of classification. *Journal of Experimental Psychology: Learning*, Memory, and Cognition, 10(1), 104–114. 10.1037/0278-7393.10.1.104

Nosofsky, R. M. (1986). Attention, similarity, and the identification-categorization relationship. Journal of Experimental Psychology. General, 115(1), 39–61. 10.1037//0096-3445.115.1.39

Nosofsky, R. M., Palmeri, T. J., & McKinley, S. C. (1994). Rule-plus-exception model of classification learning. Psychological Review, 101(1), 53.

Palmeri, T. J., & Gauthier, I. (2004). Visual object understanding. Nature Reviews Neuroscience, 5(4), Article 4. 10.1038/nrn1364

Palmeri, T. J., & Nosofsky, R. M. (1995). Recognition memory for exceptions to the category rule. *Journal of Experimental Psychology: Learning*, Memory, and Cognition, 21(3), 548.

Peirce, J., Gray, J. R., Simpson, S., MacAskill, M., Höchenberger, R., Sogo, H., Kastman, E., & Lindeløv, J. K. (2019). PsychoPy2: Experiments in behavior made easy. Behavior Research Methods, 51(1), 195–203. 10.3758/s13428-018-01193-y

Posner, M. I. (1980). Orienting of attention. The Quarterly Journal of Experimental Psychology, 32(1), 3–25. 10.1080/00335558008248231

R Core Team. (2021). R: A Language and Environment for Statistical Computing [Computer software]. R Foundation for Statistical Computing. https://www.R-project.org/

Rehder, B., & Hoffman, A. B. (2005). Eyetracking and selective attention in category learning. Cognitive Psychology, 51(1), 1–41. 10.1016/j.cogpsych.2004.11.001

Seger, C. A., Braunlich, K., Wehe, H. S., & Liu, Z. (2015). Generalization in Category Learning: The Roles of Representational and Decisional Uncertainty. Journal of Neuroscience, 35(23), 8802–8812. 10.1523/JNEUROSCI.0654-15.2015

Shepard, R. N., Hovland, C. I., & Jenkins, H. M. (1961). Learning and memorization of classifications. Psychological Monographs: General and Applied, 75(13), 1–42. 10.1037/h0093825

Smith, J. D., & Minda, J. P. (1998). Prototypes in the mist: The early epochs of category learning. *Journal of Experimental Psychology: Learning*, Memory, and Cognition, 24(6), 1411–1436. 10.1037/0278-7393.24.6.1411

Theeuwes, J. (1991). Exogenous and endogenous control of attention: The effect of visual onsets and offsets. Perception & Psychophysics, 49(1), 83–90. 10.3758/BF03211619

Theeuwes, J. (1992). Perceptual selectivity for color and form. Perception & Psychophysics, 51(6), 599–606. 10.3758/BF03211656

Turner, B. M., & Sloutsky, V. M. (2024). Cognitive Inertia: Cyclical Interactions Between Attention and Memory Shape Learning. Current Directions in Psychological Science, 09637214231217989. 10.1177/09637214231217989

van Moorselaar, D., Theeuwes, J., & Olivers, C. N. L. (2014). In competition for the attentional template: Can multiple items within visual working memory guide attention? Journal of Experimental Psychology. Human Perception and Performance, 40(4), 1450–1464. 10.1037/a0036229

Weichart, E. R., Evans, D. G., Galdo, M., Bahg, G., & Turner, B. M. (2022). Distributed Neural Systems Support Flexible Attention Updating during Category Learning. Journal of Cognitive Neuroscience, 1–19. 10.1162/jocn_a_01882

Wolfe, J. M. (2021). Guided Search 6.0: An updated model of visual search. Psychonomic Bulletin & Review, 28(4), 1060–1092. 10.3758/s13423-020-01859-9

Woodman, G. F., Luck, S. J., & Schall, J. D. (2007). The Role of Working Memory Representations in the Control of Attention. Cerebral Cortex, 17(suppl_1), i118–i124. 10.1093/cercor/bhm065

Yu, X., & Geng, J. J. (2019). The attentional template is shifted and asymmetrically sharpened by distractor context. Journal of Experimental Psychology: Human Perception and Performance, 45(3), 336–353. 10.1037/xhp0000609

Zeithamova, D., & Bowman, C. R. (2020). Generalization and the hippocampus: More than one story? Neurobiology of Learning and Memory, 175, 107317. 10.1016/j.nlm.2020.107317

Zeithamova, D., Dominick, A. L., & Preston, A. R. (2012). Hippocampal and Ventral Medial Prefrontal Activation during Retrieval-Mediated Learning Supports Novel Inference. Neuron, 75(1), 168–179. 10.1016/j.neuron.2012.05.010

Zhang, Z., & Carlisle, N. B. (2023). Explicit attentional goals unlock implicit spatial statistical learning. Journal of Experimental Psychology. General, 152(8), 2125–2137. 10.1037/xge0001368

